# Cervical Spinal Cord Injury Leads to Injury and Metabolic Dysfunction of the Lung

**DOI:** 10.1101/2021.08.16.455269

**Authors:** Emily E. Huffman, Brittany E. Dong, Harrison A. Clarke, Lyndsay E.A. Young, Derek B. Allison, Ramon C. Sun, Christopher M. Waters, Warren J. Alilain

## Abstract

High-cervical spinal cord injury (SCI) often disrupts respiratory motor pathways and disables breathing in the affected population. Moreover, cervically injured individuals are at risk for developing acute lung injury (ALI), which predicts substantial mortality rates. While the correlation between ALI and SCI has been found in the clinical setting, the field lacks an animal model to interrogate the fundamental biology of this relationship. To begin to address this gap, we performed an experimental cervical SCI and assessed lung injury in adult rats. We demonstrate that animals display signs of ALI two weeks post-SCI. We also observed aberrant N-glycan metabolism determined by matrix-assisted laser desorption/ionization mass spectrometry imaging. Collectively, we establish for the first time a model of ALI after SCI at an acute time point that can be used to monitor the progression of lung damage, as well as identify potential targets to ameliorate ALI.

## Introduction

Cervical spinal cord injuries (SCIs) can silence descending signals from the brainstem that control diaphragmatic activity. This injury can consequently impair breathing, making individuals prone to the development of profound respiratory complication ^1–3^. In fact, respiratory complication is the leading cause of morbidity and mortality for cervical SCI individuals ^4^. Two common forms of these complications found in SCI are acute lung injury (ALI) and its most severe form, acute respiratory distress syndrome (ARDS). Both complications introduce significant risks of mortality that may occur in association with a myriad of etiologies such as trauma, pneumonia, shock, aspiration, and sepsis ^5,6^. Inflammatory responses to these neutrophil mediated injuries contribute damage to the vascular endothelium and alveolar epithelium, where long-term repercussions can persist even after resolution of lung injury ^7,8^.

Recent evidence suggests that persons with an SCI hold a significant risk of developing ARDS/ALI. Most notably, those with a cervical injury hold the greatest risk and the highest mortality rate compared to other levels of SCI ^9^. This observation is not limited to SCI alone. In fact, ARDS/ALI has been found in humans following neurotrauma, including vertebral column fracture, stroke, and traumatic brain injury ^9–11^. In addition to the implications for lung damage and inhibition of ventilatory function, there is evidence that ARDS/ALI also impacts neurologic outcomes following central nervous system injury ^12^. These substantial consequences present an exigent call to characterize the peripheral repercussions of SCI. While many comorbidities and complications of SCI have gained attention in recent years, the effect of a cervical SCI on the lungs, especially the characterization of their health, function, and cellular composition, has not been explored in the animal model ^13,14^. Currently, the field lacks an animal model to interrogate the molecular underpinning of ALI after cervical SCI, which is a major barrier to identifying therapeutic windows and treatment opportunities.

In this study, we report a model of cervical SCI-induced ALI/ARDS in rats. We found classic markers of lung injury after SCI that persist two weeks after the initial time of cervical injury. We performed a thorough assessment of lung injury including severity of edema, protein infiltration in the bronchoalveolar lavage fluid, neutrophil and cellular levels, cytokine concentrations, N-linked glycans, and histopathological reports which enabled a thorough assessment of lung injury. This study demonstrates that features of lung injury ensue after experimental cervical SCI, establishing a promising model of ALI/ARDS at an acute time point that can be used to characterize and monitor the progression of lung injury after SCI, as well as provide a means to test potential therapies.

## Materials and Methods

### Animals

All procedures were in accordance with Institutional Animal Care and Use Committee (IACUC) guidelines. Thirty female Sprague Dawley retired breeder rats (12 months) from Charles River Laboratories, Inc. were housed under standard light/dark cycles with *ad libitum* access to food and water. Animals were monitored daily following surgery.

### C2 Hemisection Spinal Cord Injury

C2 Hemisection was conducted as previously described ^15^. Briefly, animals were anaesthetized with 4% isoflurane and prepared for surgery by shaving and cleaning from the ears to the shoulders with alternating bouts of Betadine and 70% ethanol. Body temperature was maintained throughout the procedure by a heating pad underneath the animal. Under aseptic conditions, a dorsal midline incision was performed to expose the cervical region of the spinal column. A C2 laminectomy allowed for visualization of the dorsal cervical spinal cord. After durotomy, a 27-gauge needle was bent and inserted into midline at C2 and dragged laterally to sever the left hemi-cord. This process was repeated for a total of three times to ensure completion. After C2 hemisection (C2Hx), the layers of musculature were sutured together and the skin was clipped closed. The animals were given Buprenorphine (0.02-0.05mg/kg), Carprofen (5mg/kg), and saline when removed from anesthesia. These post-operative care procedures were continued for three days following injury.

### Experimental Design

Fifteen rats were given a cervical SCI, while another fifteen were left intact as naïve. Sham animals were not included as the injury mimics a vertebral column fracture, which has also been shown to induce ALI/ARDS in the clinical setting, rendering it an ineffective control for studying the effects of an SCI on the lungs ^9^. The SCI and naïve animals were each randomly assigned to three groups: the edema group, BAL group, and histology/MALDI-MSI group. These 3 groups each contained 5 naïve animals and 5 SCI animals. This division allows for each group to have 10 animals total, with 20 lungs to study.

### Pre-collection Procedure and Sacrifice

Animals were deeply anesthetized under 4% isoflurane (SomnoSuite Low-Flow Anesthesia System) and subsequently overdosed by isoflurane administration 2 weeks after SCI. Rats received a laparotomy and the abdominal aorta was transected as a secondary means of sacrifice. All procedures were conducted immediately post-mortem.

### Bronchoalveolar Lavage Fluid Collection

Following sacrifice, the ribcage was reflected and suture was guided under each primary bronchus, preparing for ligation. An 18-gauge tracheal cannula connected to a 3 mL syringe filled with 2 mL of sterile, cold 0.1% EDTA was inserted without force into the left bronchus and ligated with suture. The syringe was compressed slowly, taking 1 minute to fill the lung. Insertion was verified by inflation of the lung. After a 1-minute pause, bronchoalveolar lavage (BAL) fluid was withdrawn slowly, taking 1 minute to remove from the left lung, and deposited into sterile vials on ice. This was repeated by attaching a second syringe filled with 2 mL of sterile solution to the inserted cannula. Keeping the cannula in the respiratory tract, the needle was then guided into the right primary bronchus and the procedure was repeated. In total, 4 mL of lavaged fluid were collected from each lung.

### Pulmonary Edema

Following sacrifice, the heart and lungs were pulled out of the animal *en bloc*, cutting all connective tissue and vessels in the process. Once removed, the lungs were separated, blotted until visibly dry, and placed in pre-weighed 15 mL conical vials. The wet lungs were weighed before placement in a hybridization oven at 65 degrees Celsius uncapped. After 48 hours, the desiccated lungs were weighed again. The wet/dry weight ratio was calculated by dividing the mass of the wet lung by the mass of the dry lung.

### Histology

A 16-gauge blunted cannula was inserted into the trachea and secured with suture. Sixty mL cold PBS were pushed through the right ventricle of the heart to clear the lungs of blood. The cannulated lungs were then removed from the pleural cavity along with the heart and pressure fixed (25 cm H_2_O) *en bloc* with 10% formalin. The left and right lungs were then dissected from each other and placed in tissue cassettes in 10% formalin for 24 hours. Cassettes were transferred to 70% ethanol for another 24 hours. Lungs were promptly embedded in paraffin, serially sectioned at 5 *μ*m, and mounted on slides. Injury severity was assessed by a blinded pathologist after hematoxylin and eosin staining (H&E stain).

### Cytokine Assessment

Collected BAL was centrifuged at 200 x g for 10 minutes at 4 °C and the supernatant was immediately frozen. TNF-α, IL-1β, IL-4, IL-6, and keratinocyte chemoattractant/ human growth-related oncogene (KC/GRO) concentrations were measured in the supernatant via mesoscale discovery assay (MSD), which uses electrochemiluminescence as a detection technique for higher sensitivity.

### Protein Assessment

Protein concentrations were measured via Bicinchoninic Acid assay (BCA). Briefly, 25 *μ*L of BAL supernatant were mixed with 100 *μ*L of reagent and incubated at 37 °C for 30 minutes. Protein concentration was analyzed at 562 nm and compared to a standard curve constructed using known protein concentrations.

### Cellular Assessments

Pellets from centrifuged BAL were resuspended in 500 *μ*L PBS. Cell counts were performed with Trypan blue exclusion and approximately 30,000 cells were cytospun onto microscopy slides. After drying, the slides were fixed and stained with a Diff-Quik stain. Total neutrophils were measured by morphologically identifying neutrophils per 200 cells observed.

### Matrix-Assisted Laser Desorption Ionization Mass Spectrometry Imaging

Formalin-fixed paraffin embedded lungs (FFPE) were processed as previously described ^16^. In brief, FFPE lungs were sectioned at 5 *μ*m on glass slides. Tissues were dewaxed and rehydrated followed by an antigen retrieval process in citraconic anhydride buffer. Recombinant PNGase F (0.1 *μ*g/mL) was applied by a M5 TM robotic Sprayer (HTX Technologies LLC, Chapel Hill, NC). Following incubation in a humidity chamber for 2 hours, slides were desiccated for 24hours. The following day, 7mg/mL of alpha-Cyano-4-hydroxycinnamic acid in 50% acetonitrile with 0.1% TFA were applied to each slide by the robotic sprayer. Slides were subsequently stored in a desiccator prior to MALDI-MSI analysis.

A Waters Synapt G2Si mass spectrometer (Waters Corporation, Millford, MA) equipped with a Nd:YAG UV laser with a spot size of 100 *μ*m was used to detect N-glycans at X and Y coordinates of 100 *μ*m. Following data acquisition, mass spectra were analyzed by High Definition Imaging (HDI) Software (Waters Corporation) for mass range 500-3500m/z. Three regions of interest were determined for each lung, pixel intensities were averaged and normalized by total ion current. Representative N-glycans were generated by Glycworkbench.

### Statistical Analysis

All data are presented as the mean ± SEM. All numerical data were analyzed by a two-tailed unpaired student t-test. For all analyses, P<0.05 was considered significant. Following MALDI-MSI analysis of N-gylcans, Metaboanalyst 5.0 was used to generate heatmaps and principal component analyses ^17^.

## Results

### Pulmonary Edema Developed after SCI

To determine whether there was an excess of fluid in the lungs after acute SCI, we evaluated the wet/dry ratio from the lungs of naïve and 14dpi SCI animals. The wet/dry ratio has been found to correlate with the results of other measures of lung injury ^18^. We found that C2Hx elicited a significant increase in the wet/dry ratio of the lungs of SCI animals (4.89±0.06) compared to naïve lungs (4.72±0.04) (P<0.05) (Fig. 1A). There was a trend towards a greater wet/dry ratio in the left lungs compared to the right lungs following C2Hx (N=5) (Figure 1B). However, with a small sample size, we were unable to evaluate the significance of this trend ^19^.

**Figure 1.**
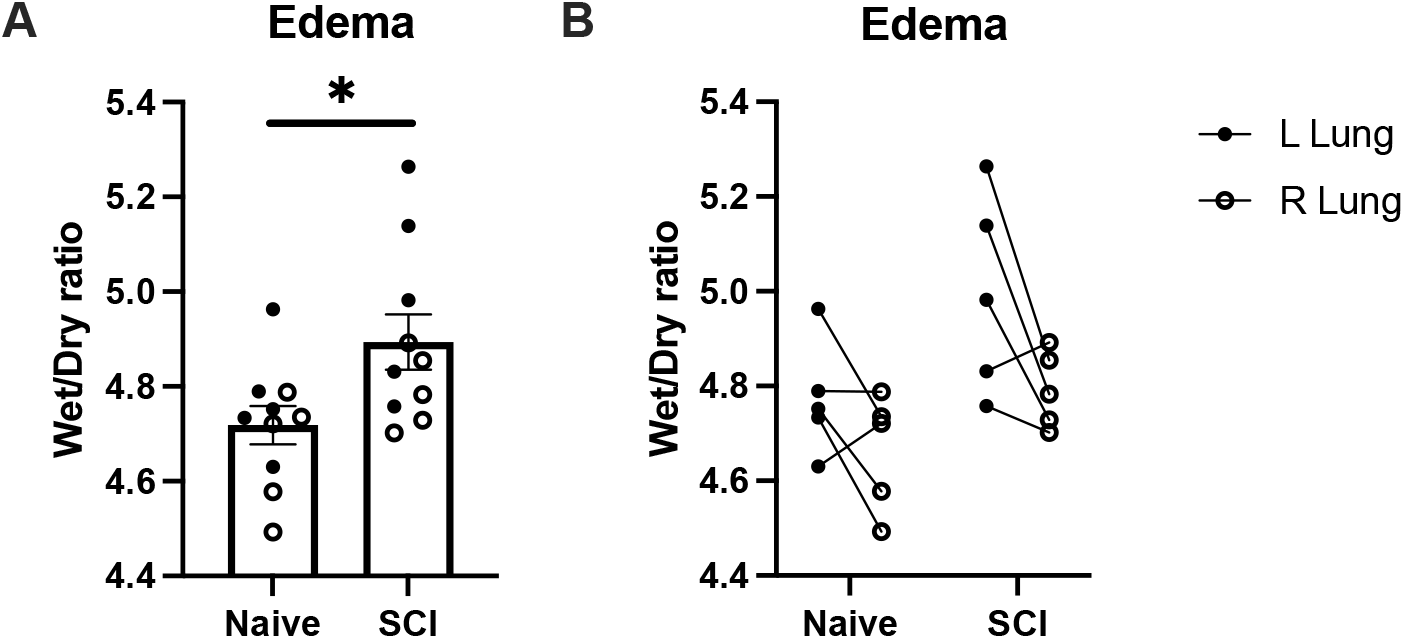
Wet/Dry ratios of excised lungs were significantly increased two weeks post-SCI compared with naïve rats (A; N=10, P<0.05). The left lung presented a trend towards a greater wet/dry ratio (B; N=5). The significance of this finding was not analyzed due to small Ns.

### SCI Increases BAL Cell Counts but Not Protein Levels in Rats

To further characterize the severity of injury, we evaluated the migration of cells into the lungs after SCI ^20^. Following SCI, the total number of cells increased significantly (1.41± 0.04 ×10^7^) compared to naïve lungs (4.31± 0.02 ×10^6^) (P<0.05). Additionally, a differential cell count revealed that the number of neutrophils was elevated in SCI animals (1.10± 0.18 ×10^5^) compared to naïve (7.79±1.29 ×10^3^) (P<0.0001) (Fig 2 A-D). Using these observations, we estimated that neutrophils comprised 1.8% of the BAL prior to injury and 8.6% post-SCI. These results show that not only did the number of cells increase in the BAL, but the ratio of neutrophils found in the lungs also increased after SCI. However, as shown in Fig. 2 E-F, protein levels in the BAL were not significantly increased in the SCI rats (0.133±0.006 μg/μl) compared to naïve (0.124±0.005 μg/μl) (Fig. 2 E-F). Again, there appeared to be a trend towards the left lung having both a greater cell count and a larger neutrophil count compared to the right lung (N=5).

**Figure 2.**
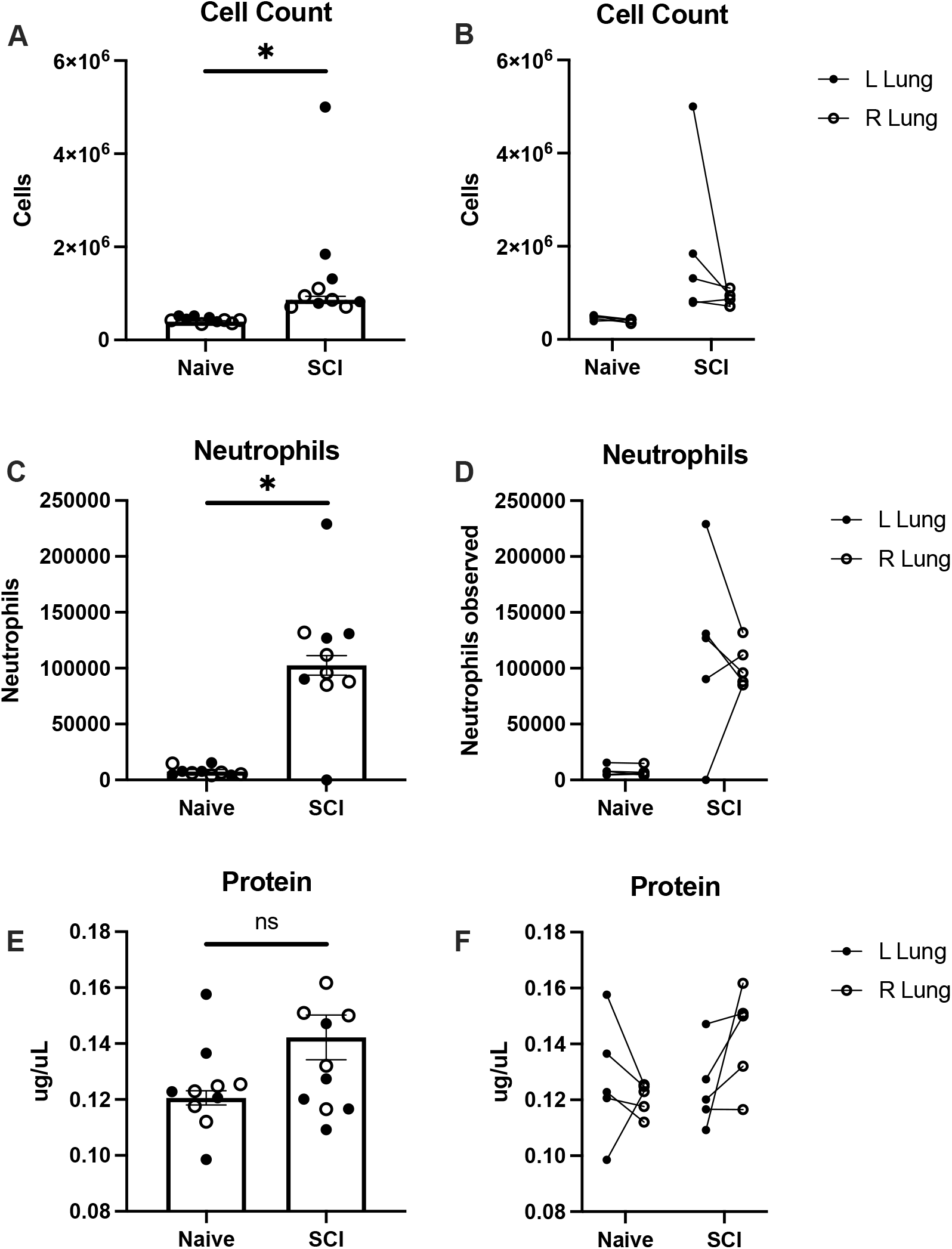
Changes in cell numbers and neutrophil counts were significantly increased two-weeks post-SCI compared with naïve rats. Measurements of total cell count (A-B), neutrophils (C-D), and protein concentration (E-F) in bronchoalveolar lavage fluid (BAL) from naïve and two weeks post-SCI rats. There was a significant increase in total cell count and neutrophils following SCI (A, C; N=10, P<0.05), but no detected change in protein (E N=10, P>0.05). The left lung trends towards a greater cell count and neutrophil levels following SCI (B, D; N=5). However, the significance of this finding was not analyzed due to small Ns.

### SCI Modulate Cytokine Levels in BAL

To further investigate the increase in cells found in the BAL after SCI, we measured levels of the neutrophil chemoattractant KC/GRO, as well as other indicators of inflammation including the cytokines TNF-α, IL-1β, and IL-6, whose concentrations have been observed to increase after lung injury ^21–23^. MSD results showed significantly increased levels of KC/GRO (100.0 ±12.33 μg/μl) and TNF-α (0.50±0.06 μg/μl) in SCI animals compared to naïve (64.04±2.28 μg/μl), (0.31±0.03 μg/μl) respectively (Fig. 3 A-D). IL-1β levels remained unchanged two weeks after injury (3.06±0.41) compared to naïve (2.23±0.25 μg/μl) (Fig E-F).

**Figure 3.**
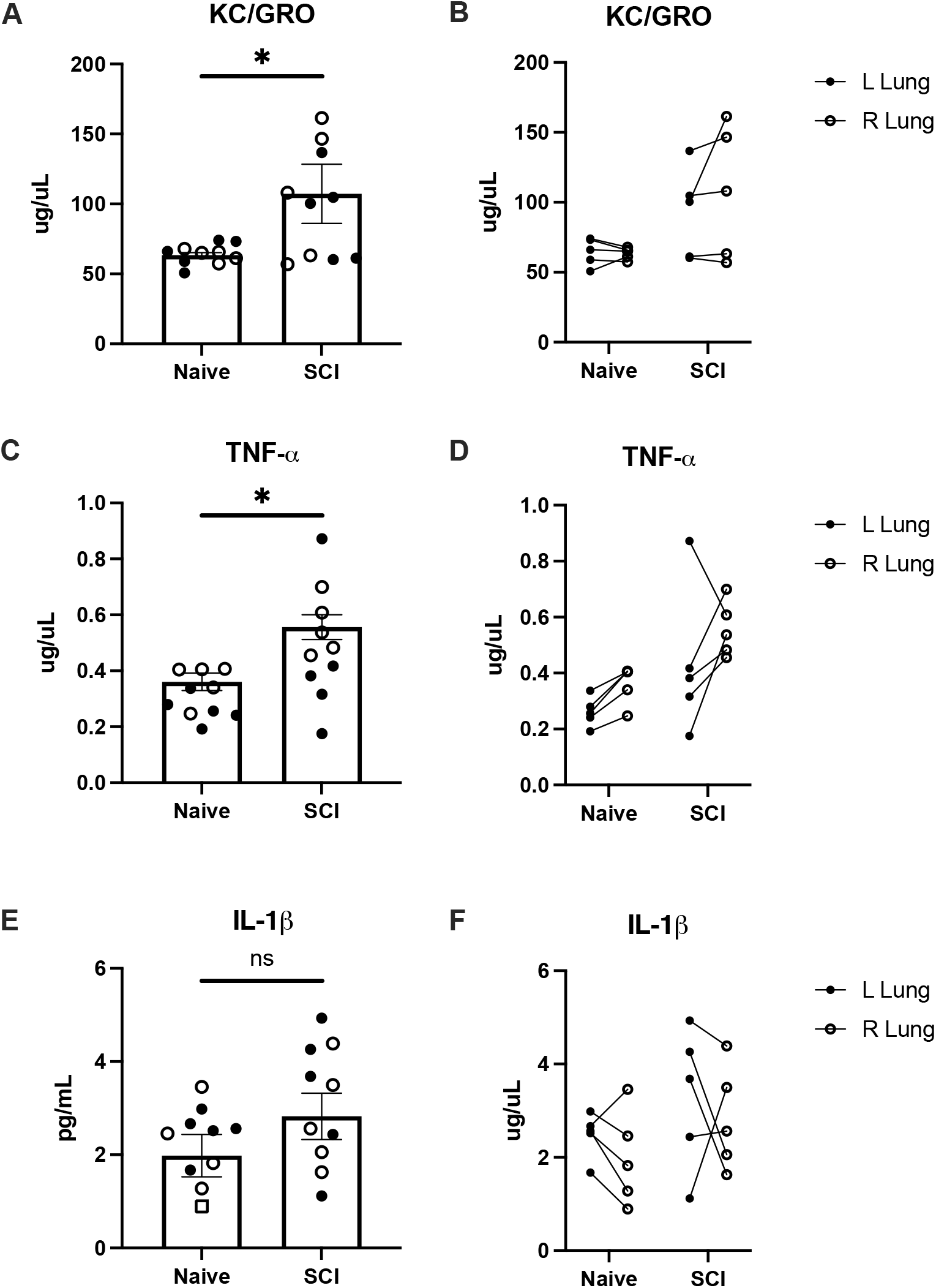
Levels of KC/GRO and TNF- α were significantly increased two-weeks post-SCI compared with naïve rats. Measurements of KC/GRO (A-B), TNF-α (C-D), and IL-1β (E-F) in bronchoalveolar lavage fluid (BAL) from naïve and two weeks-post SCI rats. There was a significant increase in levels of KC/GRO and TNF-α following SCI (A, C; N=10, P<0.05), however, there was no significant change in levels of IL-1β following SCI. Side-specific differences were not found following SCI due to small Ns (B, D, F; N=5).

**Figure 4.**
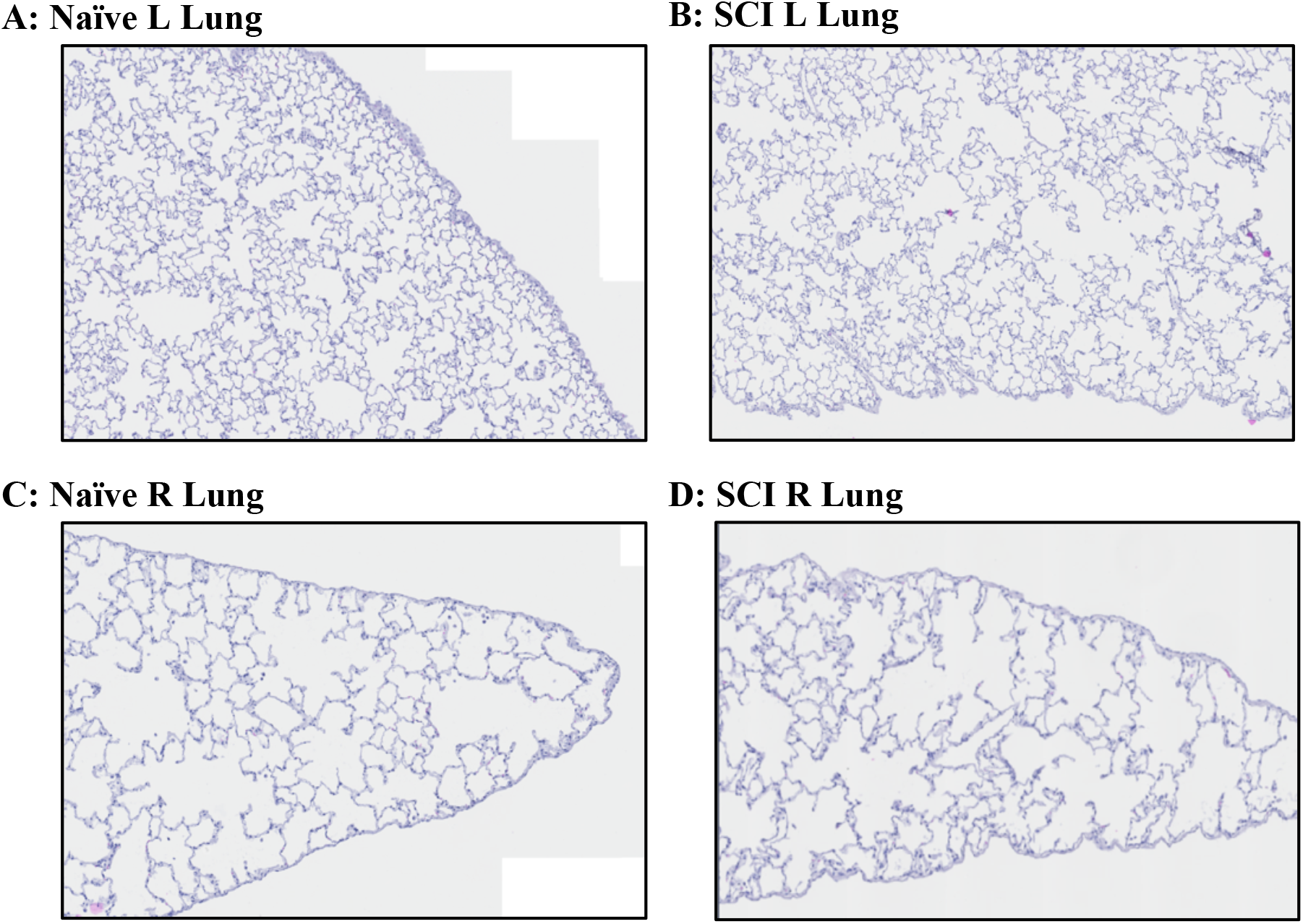
No histopathologic difference were observed in the lungs from naïve and two weeks post-SCI rats. Lungs were essentially normal (H&E stain, low power). (A: representative naïve left lung; N=5) (B: representative SCI left lung with scattered parabronchial lymphoid aggregates; N=5) (C: representative naïve right lung; N=5) (D: representative SCI right lung; N=5).

Levels of IL-6 were under the range of detection in 15 out of 20 samples. However, the in-range samples belonged to SCI animals.

### No Histopathologic Changes Were Observed in the Lungs After Acute SCI

Histological analysis by a blinded pathologist using H&E staining revealed similar findings in both SCI and naïve animals. More specifically, both groups had rare scattered small foci containing foamy pulmonary alveolar macrophages, mild chronic interstitial inflammation, and mild interstitial fibrosis. However, the majority of the examined lung parenchyma was relatively normal in appearance in all cases. With a more severe acute lung injury, we would expect to see an accumulation of intra-alveolar immune infiltrates, alongside more robust and diffuse interstitial inflammatory infiltrate and microvascular congestion. Lack of histopathology features suggest we are at early stages of lung injury, thereby establishing the treatment window for the future testing of therapeutic options.

### SCI Is Associated With Changes in N-linked Glycosylation of the Lung

In order to determine whether SCI caused metabolic changes in lung tissue, we performed crude carbohydrate analysis of the lung using periodic acid-schiff (PAS) stain that revealed broad changes in complex carbohydrates ^24–26^. Similar to H&E analysis, PAS staining showed no detectable changes between naïve and ALI following SCI (Fig. 5C and 6C). To further investigate whether N-linked glycan changes occur in ALI, we performed *in situ* analysis of N-linked glycomics by enzyme-assisted matrix-assisted laser desorption/ionization (MALDI) mass spectrometry imaging. First, we performed multiple multivariate analysis that include supervised clustering heatmap and partial least squares discriminant analysis (PLS-DA) on both left and right lung with ALI (Fig. 5A-B & 6A-B). Both clustering heatmap and PLS-DA analysis uniquely separated naïve and SCI injured lungs (Fig. 5A-B & 6A-B). Interestingly, variable importance in projection (VIP) values from PLS-DA reveal heterogenous changes in N-link glycans between left and right lungs (Fig. 5B & 6B). Finally, we performed gross tissue structural analysis using the MALDI dataset and found that changes in glycans are consistent across entire lung regions and not confined to specific regions (Fig. 5D & 6D). Collectively, MALDI imaging analysis of N-linked glycomics suggest distinct aberrant N-glycan metabolism is associated with ALI after SCI and these changes are lobe dependent. Further, spatial analysis showing universal changes across entire lobe suggests the possibility of mechanical defects after SCI rather than localized cellular damage.

**Figure 5.**
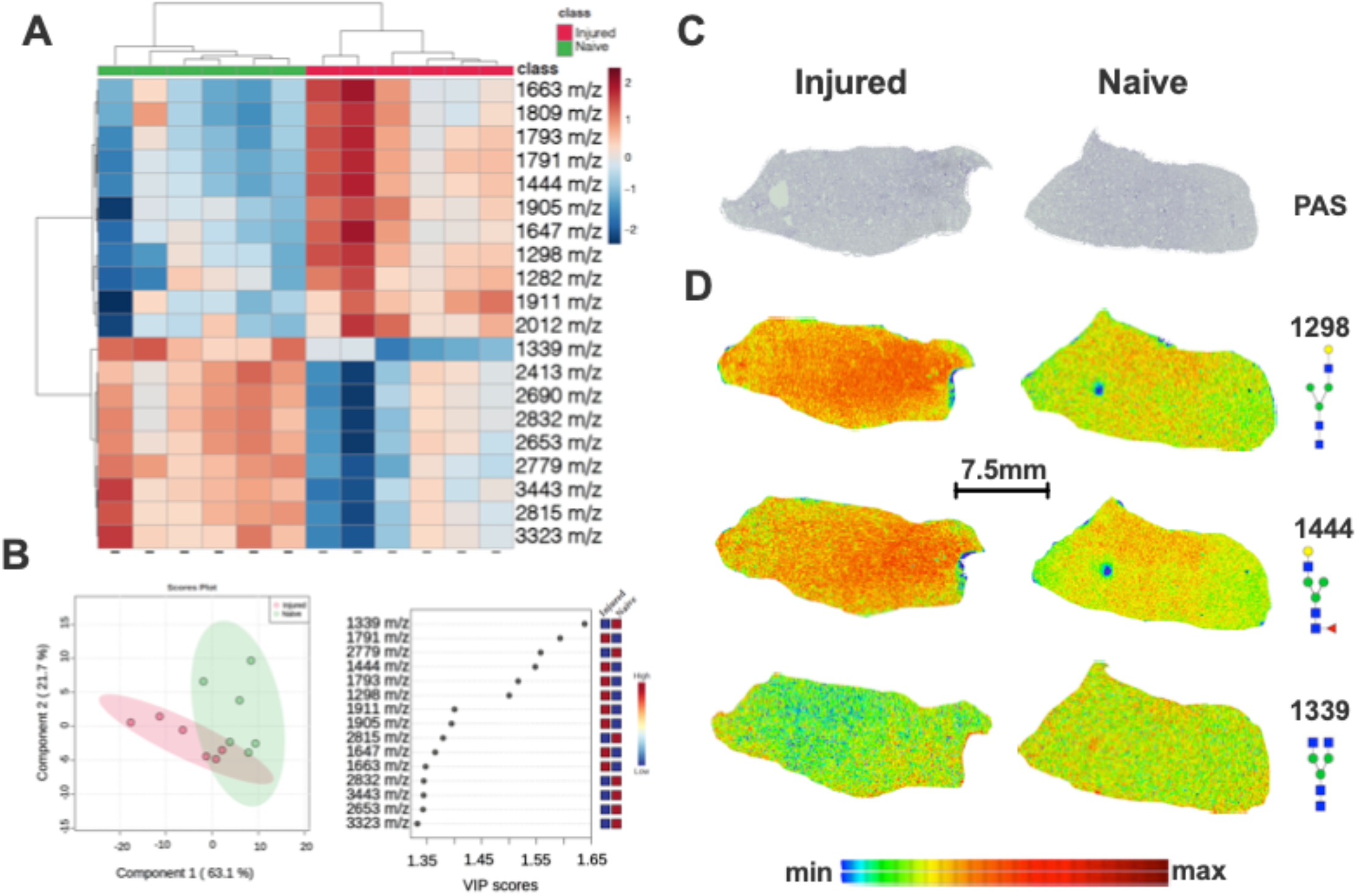
MALDI mass spectrometry imaging analysis of the left lung following SCI (N=5). Multivariant clustering heat analysis shows separation of SCI and naïve lungs using top 25 most changed N-linked glycans (A). (B left panel) PLS-Da analysis shows 2D clustering of SCI and naïve rat lungs (B left panel). VIP scores of most changed N-glycans in whole lung (B right panel). There was no significant PAS difference the left lungs from SCI and naïve rats (C; N=5). Representative images of selected N-glycans and their spatial distribution across the whole lung (D; N=5).

**Figure 6.**
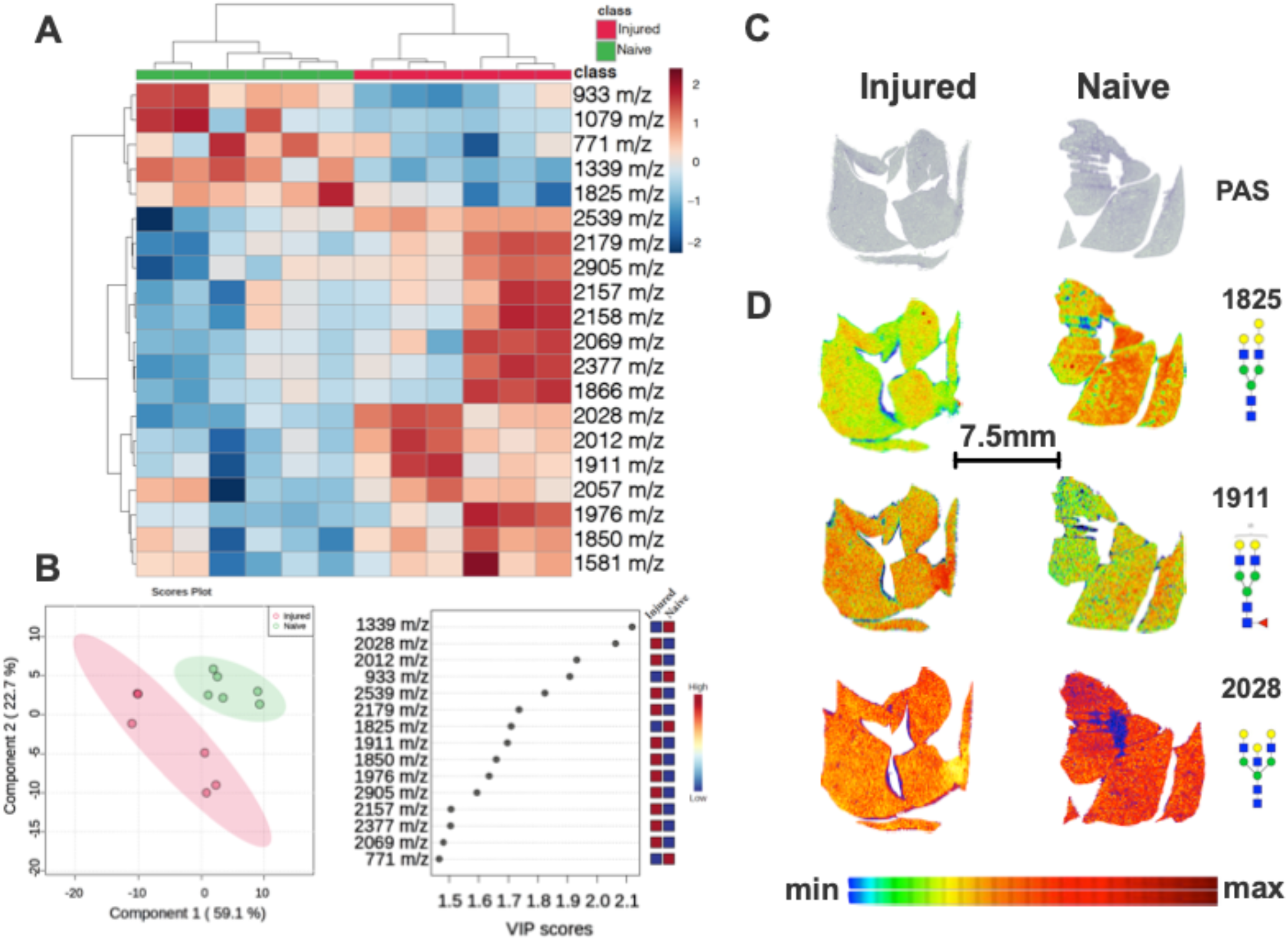
MALDI mass spectrometry imaging analysis of the right lung following SCI (N=5). Multivariant clustering heat analysis shows separation of SCI and naïve lungs using top 25 most changed N-linked glycans (A). (B left panel) PLS-Da analysis shows 2D clustering of SCI and naïve rat lungs (B left panel). VIP scores of most changed N-glycans in whole lung (B right panel). There was no significant PAS difference the right lungs from SCI and naïve rats (C; N=5). Representative images of selected N-glycans and their spatial distribution across the whole lung (D; N=5).

## Discussion

A majority of individuals with SCI develop secondary complications within the first two weeks of injury ^27^. Specifically, individuals with cervical SCI have a high incidence of respiratory complications and can develop ARDS/ALI after injury ^9,28^. We investigated this clinical observation in the rat model in an effort to capture the effects of SCI on the lungs. Overall, this study demonstrates that mild lung injury does occur following cervical SCI. We observed signs of mild ARDS/ALI at two weeks after cervical SCI with trends towards greater injury in the lung ipsilateral to SCI. These findings establish a model of ALI/ARDS at an acute time point that can be used to characterize and monitor the progression of lung injury after SCI, as well as a means to investigate potential therapeutic strategies.

ARDS/ALI is a progressive disease with various forms of damage occurring at different timepoints depending upon the initiating event. The temporal developments of lung injury following SCI have not been established in the rat model ^29^. However, our data appear to indicate the exudative stage of ALI, with evidence of edema and infiltration of cells into the alveoli without indications of fibrosis. These observations two weeks after SCI suggest a disruption of the epithelial-endothelial barrier in the alveoli.

ARDS/ALI characteristically results in the increased permeability of the alveolar-capillary barrier. The damage of this barrier facilitates the influx of fluid into the alveoli. We assessed edema by calculating the wet/dry ratio for each lung and we found that animals with SCI had increased edematous fluid in both the left and right lungs compared to uninjured controls. Interestingly, despite utilizing a unilateral SCI model, there was no significant difference in edema between the left and the right lung, suggesting that our model of SCI did not specifically target either lung. However, there is a reasonably strong trend that the ipsilateral lung had a greater wet/dry ratio compared to the contralateral lung in SCI animals. With a small sample size, we were unable to evaluate the significance of this trend ^19^. Another indicator of lung injury is the recruitment of immune cells into the airspace of the lungs. As expected, we observed primarily alveolar macrophages in the cell counts of uninjured animals. After SCI, the population of cells in the alveoli increased primarily due to the recruitment of neutrophils. Because we observed a substantial increase in both total cell number and neutrophils in the BAL after SCI, we evaluated KC/GRO as it is known to mediate neutrophil recruitment and activation ^30^. We found an increase in KC/GRO in the BAL following SCI. We demonstrated that the increase in neutrophils and KC/GRO persisted for 14 days following SCI. KC/GRO has functional homology with IL-8 with chemotactic and proinflammatory activity in the rodent ^31^. High levels of KC/GRO in the BAL from ARDS/ALI patients are associated with increased neutrophils in the injured lungs ^32^.

Knowing that KC/GRO levels were elevated, we assayed cytokines involved in its inflammatory cascade ^33^. We observed increased levels of TNF-α in the collected BAL fluid from SCI rats. TNF-α has been proposed as a mediator of lung injury and its neutralization ameliorates the injury ^21,22^. Interestingly, there were no significant changes in IL-1β levels after SCI and IL-6 levels were undetectable in naïve and SCI animals. The inability to detect changes in IL-1β and IL-6 may be due to these cytokines’ varying temporal profiles, with IL-6 peaking hours after induction of lung injury in the mouse model, for example ^17^. However, more timepoints need to be evaluated after SCI to provide further insights into the progression of lung injury following SCI.

Increased protein concentration in the BAL is another indicator of lung injury and vascular permeability in both rodents and humans ^34^. Damage to the alveolar-capillary barrier allows protein from the blood to enter the lungs at the level of the capillaries ^35^. However, despite increased wet/dry ratios in SCI rats, we did not find substantial increases in protein concentration in the BAL after SCI. One potential explanation for this finding is that fluid accumulation may have occurred due to increased hydrostatic pressure rather than loss of the alveolar-capillary barrier function. However, we do not have direct evidence for this since we did not measure pulmonary vascular pressures in this model. Our injury model is one of an indirect induction of lung injury, consistent with other forms of trauma ^36^. With the paralysis of the left hemidiaphragm as opposed to the entire diaphragm, the injury is not as severe as a complete transection or contusion SCI.

After 14 days, SCI rats showed no histopathologic differences compared to the control rats. The vast majority of the examined lungs were histologically unremarkable with both groups showing rare foci of non-specific findings. However, the presence of these limited findings in both the SCI rats and controls suggests that they are unrelated to the induced SCI. Additionally, the lack of prominent indications of diffuse alveolar damage is consistent with the lack of increased protein in the BAL. These results were expected in the acute setting of SCI, as there has likely not yet been a long enough time-course to result in significant pathologic findings. The changes that we found in Figures 1-3 may manifest into histopathological damage in more chronic timepoints following SCI.

Although we did not observe significant evidence of histological lung injury, we investigated changes in N-linked glycan metabolism as a potential early indicator of the development of lung injury. N-linked glycan biosynthesis is an understudied facet of glucose metabolism. In the lung, N-linked glycans are crucial for the differentiation of bronchoalveolar stem and alveolar type 2 cells to form the alveolar and bronchiolar lining ^37–39^. Further, N-linked glycans are critical components of mucins and surfactant proteins that maintain the liquid-air interface, reduce surface tension, and provide lubrication for the mechanical action of the lung ^40,41^. Collectively, MALDI imaging analysis of N-linked glycomics suggests distinct aberrant N-glycan metabolism is associated with ALI after SCI and that these changes are lobe dependent. Further, spatial analysis shows universal changes across the entire left and right lobes. These findings support the possibility of widespread changes in metabolism after SCI, that could potentially impact lung progenitor cell differentiation, oxygen/CO2 exchange, and movement of the lung. With these important implications in mind, aberrant N-linked glycan phenotype in our ALI model warrant further investigation.

ALI/ARDS is often induced in animal models with a direct inflammatory insult to the lungs ^42^. However, ALI/ARDS can also manifest as a result of indirect insults such as sepsis, burns, and trauma via inflammatory mechanisms ^36, 43^. After SCI, there is an intense systemic inflammatory response that affects whole organ systems ^44, 45^. Lung damage has been found in rodent models after thoracic-level SCIs with evidence for systemic inflammation as the cause ^13,46^. However, lung injury after cervical SCI cannot be solely attributed to inflammation. Individuals with cervical SCIs are at higher risk for the development of ARDS/ALI and have more severe ARDS/ALI than at any other injury level ^9^. This severity is likely because an injury to the cervical level of the spinal cord affects breathing function while simultaneously generating a general systemic inflammatory response ^47^. Our results suggest this explanation may be the case, as we demonstrate that the lung ipsilateral to the SCI trends towards more severe signs of ARDS/ALI. However, the significance of these differences was immeasurable due to small sample size ^19^. This observation may be a result of the left C2Hx injury model used in this study. In this model, the innervating signal to the left hemidiaphragm is severed above the level of the phrenic motor nucleus. This injury results in paralysis of the left hemidiaphragm, leaving the operation of the right hemidiaphragm intact. The desuetude of the left hemidiaphragm affects the pressure-differential in the left pleural cavity and lowers the tidal volume of injured animals ^15^. With a reduced capacity of the lungs to perform gas-exchange, ALI/ARDS can ensue ^48^. Future experiments will be directed toward elucidating the side-specific differences in the lungs following unilateral SCI.

We provide a means to investigate potential therapeutic strategies to ameliorate the onset and progression of lung injury after SCI in the rat model. Based on our collective findings, it would be pertinent to look to the aberrant metabolic profile of the lungs following SCI. Specifically, advances are being made in glycan-based immunotherapeutics in the treatment of breast and lung cancer ^49^. A broader use of these therapeutics may one day be applied to lung injury after SCI to target the changes in metabolism we observe in the lungs following SCI.

In summary, our studies demonstrated that animals with a cervical SCI display signs of mild ALI/ARDS including, but not limited to marked increases in alveolar neutrophil counts, heightened levels of proinflammatory cytokines and neutrophil-activating KC/GRO, intra-alveolar edema, and changes in N-glycan metabolism 14 days after injury. Collectively, this study establishes an SCI-induced ALI model that can be developed further and used to identify potential targets to ameliorate respiratory distress and lung injury after SCI.

## Acknowledgements

We thank Jessica Newton, M.S., Chris Calulot, B.S., Daimen Britch, B.S., and Lydia Strattan, Ph.D. for their assistance in completing experiments. We also thank Lydia Strattan, Ph.D., Jessica Newton, M.S., and Aaron Silverstein, B.S., B.S., for their assistance revising this manuscript.

